# Time-resolved analysis of transcription kinetics in single live mammalian cells

**DOI:** 10.1101/2021.12.23.474066

**Authors:** Hongyoung Choi, Byung Hun Lee, Hye Yoon Park

## Abstract

In eukaryotic cells, RNA polymerase II synthesizes mRNA in three stages, initiation, elongation, and termination, and numerous factors determine how quickly a gene is transcribed to produce mRNA molecules through these steps. However, there are few techniques available to measure the rate of each step in living cells, which prevents a better understanding of transcriptional regulation. Here, we present a quantitative analysis method to extract kinetic rates of transcription from time-lapse imaging data of fluorescently labeled mRNA in live cells. Using embryonic fibroblasts cultured from two knock-in mouse models, we monitored transcription of β-actin and Arc mRNA labeled with MS2 and PP7 stem–loop systems, respectively. After inhibiting transcription initiation, we measured the elongation rate and the termination time by fitting the time trace of transcription intensity with a mathematical model function. We validated our results by comparing them with steady-state fluctuation analysis and stochastic simulations. This live-cell transcription analysis method will be useful for studying the regulation of elongation and termination steps and may provide insight into the diverse mechanisms of transcriptional processes.

## Introduction

Complex regulation of eukaryotic gene expression starts with transcription, through which information in DNA is copied into mRNA. At the site of transcription, RNA polymerase II (Pol II) reads a DNA code and synthesizes mRNA, and three distinct steps can be considered to comprise the transcription procedure: initiation, elongation, and termination. Initiation is the first step in which Pol II binds to a promoter sequence at the 5ʹ end of a gene^1^. In the elongation step, Pol II reads DNA, synthesizes mRNA, and moves from the 5ʹ end to the 3ʹ end^2^. Termination is the last step in which mRNA and Pol II are released from the 3ʹ end of the gene^3,4^. Thus, three kinetic rates corresponding to each step can be defined: the initiation rate *c* (the number of Pol II binding per unit time), the elongation rate *k* (the number of base pairs transcribed per unit time), and the termination time *T_t_* (the time for mRNA separated from DNA after reaching the transcription end site). Despite many efforts to understand how transcription is regulated, limitations and challenges in quantitative measurement of these three kinetic rates still remain.

In recent years, various RNA sequencing methods have been developed to study elongation and pausing of Pol II at the promoter proximal region, intron-exon junctions, and nucleosomes^5–7^. These methods revealed not only the nascent RNA density in the steady state but also the elongation rate in time-resolved experiments. For such time-resolved measurements, transcription was induced or inhibited to track progression of Pol II after drug treatment, whereby the progression speed of the sequence read density directly reflected the elongation rate^8–11^. Nevertheless, population studies cannot resolve the transcriptional processes occurring at a single gene locus and lack the temporal resolution to investigate fast dynamics in real time.

Live-cell imaging has been applied to study transcriptional dynamics in single cells by labeling mRNA, Pol II, or transcription factors with fluorescent proteins^12,13^. Fluorescence recovery after photobleaching (FRAP) technique has been used to measure transcriptional kinetic rates by fitting recovery curves with various model functions: half recovery time^14^, linear and exponential fitting^15–17^, and solutions of rate equations^18–21^. However, there is concern about the accuracy of kinetic modeling using FRAP because recovery curves can be affected by photobleaching, diffusion, and binding of fluorescent particles; additionally, different kinetic models can fit the same experimental data almost equally well^22^. More recently, fluctuation correlation analysis of fluorescently labeled mRNA was used to measure the transcription initiation rate and dwell time of a transcript of a yeast gene^23^. In this analysis, the dwell time consists of the elongation and termination times, which cannot be separated in the steady state. Nevertheless, the authors were able to calculate the elongation rate and the termination time by comparing results from two constructs bearing a PP7 RNA stem–loop cassette in either the 5ʹ untranslated region (UTR) or 3ʹ UTR^23^. Likewise, another study measured the elongation rate of a reporter gene in living *Drosophila* embryos by comparing the results from two different positions of an MS2 stem-loop cassette^24^. After that, Coulon *et al*. developed a dual-color fluctuation correlation analysis for human β-globin reporter mRNA labeled with a PP7 cassette in the second intron and an MS2 cassette in the 3ʹ UTR^25^. Using two autocorrelation functions and a single cross-correlation function, the authors measured the kinetics of transcription and splicing of single mRNAs. Liu *et al*. also used a two-color MS2/PP7 reporter to infer the initiation rate, elongation rate and termination time of the *hunchback* gene in developing fruit fly embryos^26^. However, it would be challenging to insert ~1.5-kb-long MS2 and PP7 cassettes into two different locations in an endogenous gene, which limits the general application of the technique in mammalian cells and tissues.

In this study, we developed a time-resolved transcriptional analysis method to measure the elongation rate and termination time of an endogenous mRNA labeled with one stem–loop cassette. Previously, two knock-in (KI) mouse models, Actb-MBS^27^ and Arc-PBS^28^, were generated to label β-actin (Actb) and Arc mRNA, respectively. In the former, 24 repeats of the MS2 binding site (MBS) were inserted in the 3ʹ UTR of the Actb gene^27^ (Fig. 1A); in the latter, 24 repeats of the PP7 binding site (PBS) were knocked in the 3ʹ UTR of the Arc gene^28^ (Fig. 1B). Using mouse embryonic fibroblasts (MEFs) derived from these two mouse models, we investigated the transcriptional dynamics of the Actb and Arc genes. First, by applying the steady-state autocorrelation analysis method^23^, we measured the initiation rate and the total dwell time of each mRNA, and we then measured the elongation rate and the termination time using our time-resolved analysis method. Although we found a highly heterogeneous distribution of the kinetic parameters for each gene, there was a significant difference between Actb and Arc transcriptional kinetic rates. This approach allows quantitative measurement of all three kinetic rates of single-color labeled mRNA transcription in live cells, providing opportunities to study diverse transcriptional regulation at the single-allele level with high spatiotemporal resolution.

**Fig. 1.**
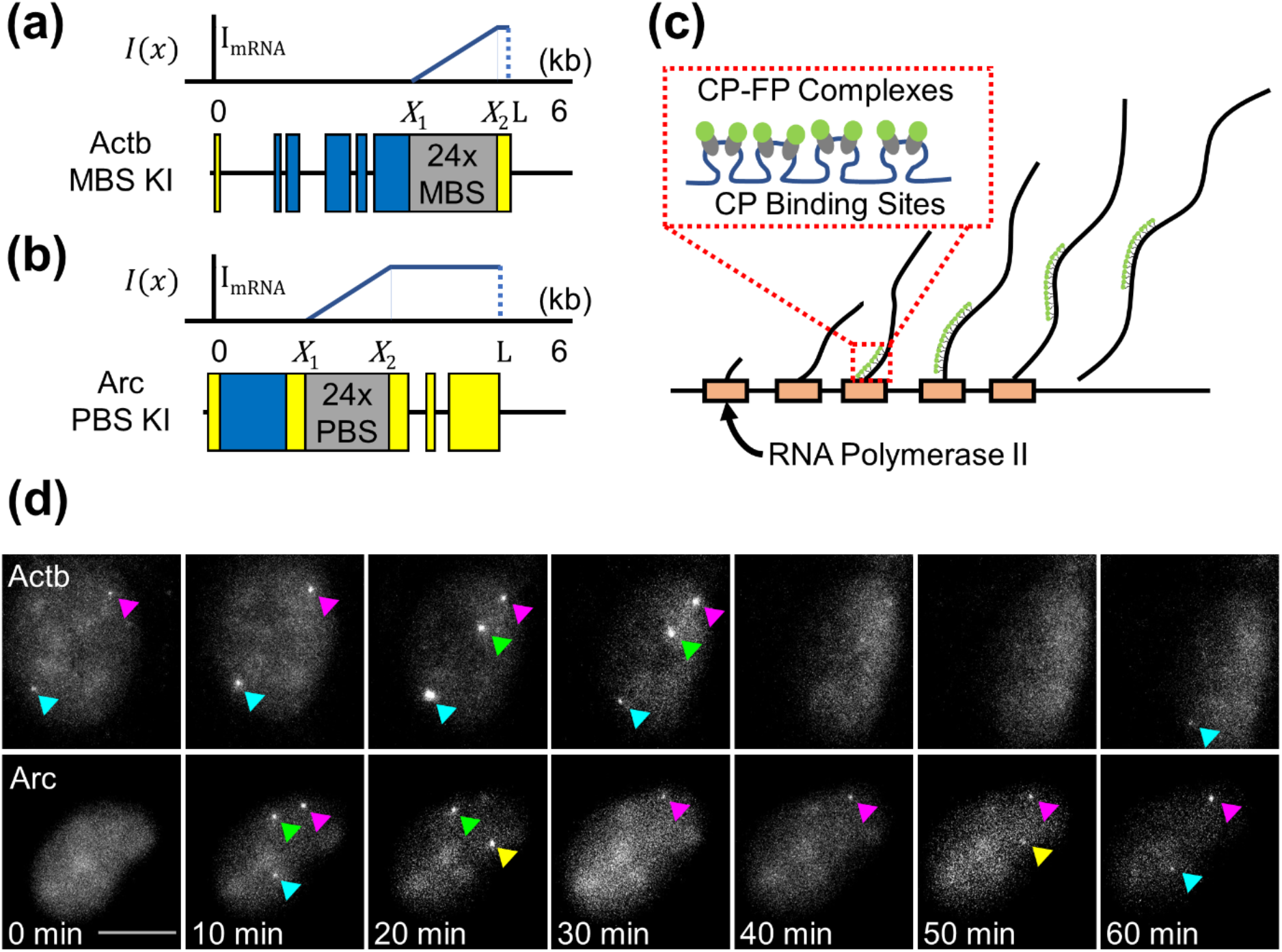
Imaging transcription using MS2 and PP7 stem–loop systems. (**a, b**) Schematic of the Actb-MBS (a) and Arc-PBS (b) knock-in constructs. The fluorescence intensity of a nascent mRNA increases as Pol II transcribes the binding sites from *X*_1_ to *X*_2_. In the post stem–loop region from *X*_2_ to *L*, the intensity remains constant (gray: binding sites, blue: coding sequence, yellow: untranslated region). (**c**) Schematic of nascent mRNAs at a transcription site. Multiple capsid proteins fused with fluorescent protein (CP-FP) complexes are bound to a single nascent mRNA. (**d**) Maximum z-projected time-lapse images of serum-stimulated MEFs. Transcription sites are observed as bright points (arrowheads) in the nucleus. Scale bar, 10 μm.

## Mathematical models

### Steady-state fluctuation correlation analysis model

Larson *et al*. introduced a mathematical model to analyze transcriptional dynamics using an autocorrelation function of a transcription site intensity trace^23^. We rederived the steady-state autocorrelation function in a more concise manner and applied the model to Actb-MBS (Fig. 1a) and Arc-PBS (Fig. 1b) mRNAs. The autocorrelation function is defined as

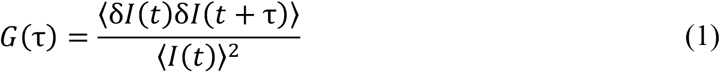

where δ*I*(*t*) = *I*(*t*) − ⟨*I*(*t*)⟩ and *I*(*t*) is the intensity of a transcription site. The bracket denotes the average intensity over time. The experimental data can be fit with the following mathematical model:

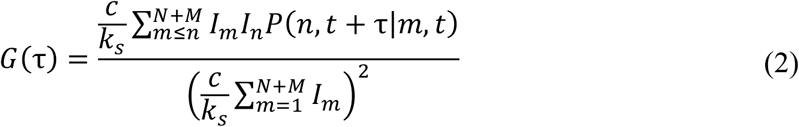

where

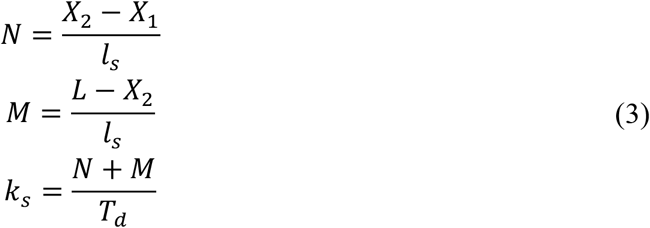

*N* and *M* are the length of the stem–loop [*X*_1_, *X*_2_] and the post stem–loop [*X*_2_, *L*] region scaled with the stem–loop size *l_s_*. *k_s_* is the effective transition rate of Pol II. *T*_d_ is the total dwell time, which includes the elongation time through the fluorescent region (the stem–loop and the post stem–loop region) and the termination time. *I_n_* is the fluorescence intensity of a pre-mRNA after Pol II transcribes the *n*th stem–loop from the 5ʹ end of the stem–loop region. This pre-mRNA intensity is described as follows:

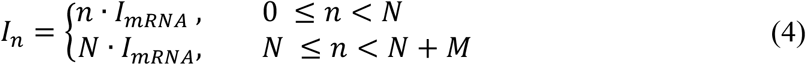

*P*(*n*, *t* + τ|*m*, *t*) in Eq. (2) is the conditional probability that a Pol II is at position *n* at time (*t* + τ) given that it has been at position *m* at time *t*.

Larson *et al*. approximated the autocorrelation function in Eq. (2) to consider two special cases in which the position of the stem–loop region was at either the 5ʹ or 3ʹ end of an mRNA. Such approximation cannot be applied for Actb-MBS and Arc-PBS mRNAs; the full autocorrelation function in Eq. (2) needs to be used for a general position of the stem–loop region. The steady-state autocorrelation function was calculated based on three assumptions: (i) initiation occurs randomly and evenly; (ii) Pol II elongates only in one direction from the 5ʹ end to the 3ʹ end, and there is no reverse translocation; and (iii) pre-mRNA is released immediately at the 3ʹ end of the gene.

With the assumption that Pol II can progress only in one direction, the probability density of Pol II dwell time *t_n_* in an arbitrary position *n* can be calculated for the irreversible process.

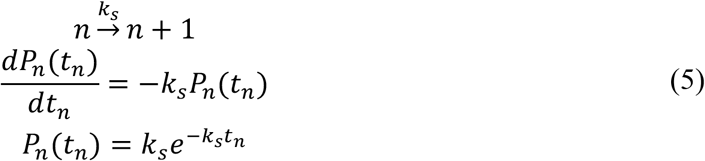

The time spent for the transition to the next position is exponentially distributed. If *t_i_* is the time spent for the transition from the *i* th stem–loop to the (*i* + 1) th stem–loop, the conditional probability *P*(*n*, *t* + *τ*|*m*, *t*) becomes

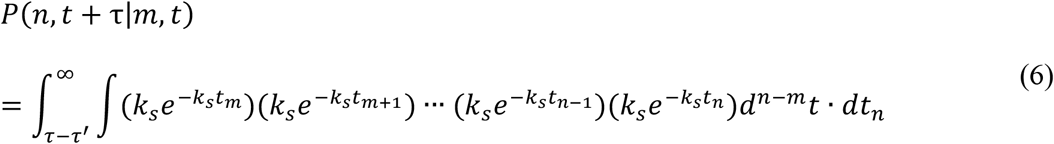

while satisfying constraints given as

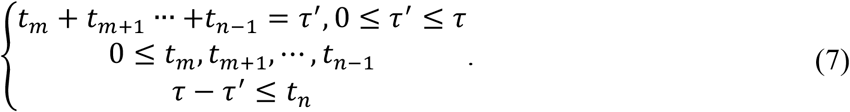

The (*n* − *m*) dimensional integral denoted as *d*^*n*−*m*^*t* is *dt_m_* ⋯*dt*_*n*−1_. Each exponential term represents the probability density that Pol II spends time *t_i_* at position *i*. Because Pol II is in the *n*th stem–loop at time (*t* + τ), if Pol II spends time τʹ from the *m*th stem–loop to the (*n* − 1)th stem–loop, the possible dwell time in the *n*th stem–loop is from *τ* − *τ*ʹ to ∞. Then, the conditional probability is described as

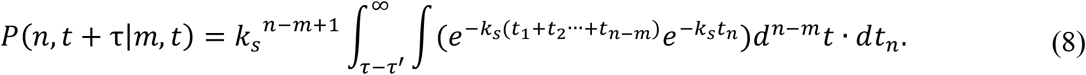

By integrating with *dt_n_* first,

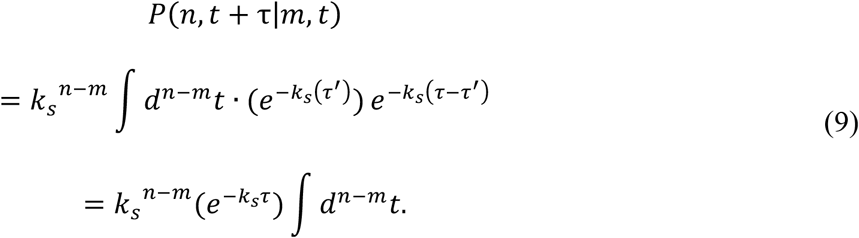

The last integral term is the (*n* − *m*) dimensional volume in the domain denoted in Eq. (7). This domain is a (*n* − *m*)-dimensional hyper pyramid in which (*n* − *m*) segments are perpendicular to each other, and their lengths are τ. The last integral in Eq. (9) is τ^*n*−*m*^/(*n* − *m*)!. Finally, the conditional probability becomes

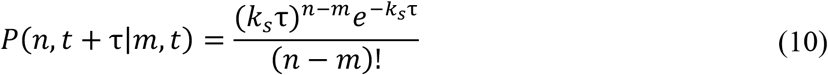

This is the Poisson distribution; *k_s_*τ is the expected number of stem–loops transcribed during time τ. The steady-state autocorrelation function can be calculated by substituting Eq. (10) into Eq. (2). The initiation rate *c* and the effective transition rate *k_s_* can be obtained by fitting the autocorrelation curve of the experimental data with Eq. (2). Using the *k_s_*value and Eq. (3), we can calculate the total dwell time *T_d_*, which includes the elongation time through the fluorescent region (the stem–loop and the post stem–loop region) and the termination time *T_t_*. Although we assumed in our derivation that pre-mRNA is released immediately at the 3ʹ end of the gene, the total dwell time calculated from *k_s_* effectively includes the termination time.

### Time-resolved transcription analysis model

We developed a time-resolved transcription analysis model to determine the transcription elongation rate *k* and the termination time *T_t_*. The dynamics of translation have previously been analyzed using the SunTag system^29^ with a mathematical model describing the decreasing fluorescence intensity of a translation site after inhibiting translation initiation^30^. Because RNA stem–loop systems are analogous to the SunTag system, we adapted this model for application to transcriptional dynamics. Although translation termination time was not considered in the previous model, we included it and derived an analytical function for the decreasing fluorescence intensity after inhibition of transcription initiation.

We assumed that Pol IIs are distributed uniformly on the gene and move toward the 3ʹ end at a constant speed *k*. For simplicity, we modeled transcription termination as a first-order irreversible process with a rate constant *a*. Hence, the number of Pol IIs residing at the 3ʹ end of the gene at time *t* and *N*(*t*) can be calculated as follows.

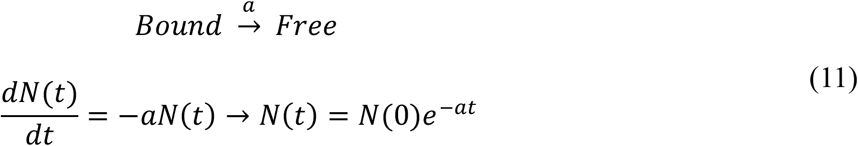

The transcription termination time was defined as *T_t_* = 1/*a*. Suppose that there is no Pol II at the 3ʹ end and that the first Pol II just reaches the end at time *t* = 0. With the transcription initiation rate *c*, the number of Pol II reaching the 3ʹ end of the gene during time interval Δ*t* is *c*Δ*t*. After another time interval Δ*t*, the number of Pol II accumulated at the 3ʹ end becomes

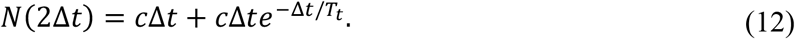

If we assume that Pol II and nascent mRNA fall off at the same time from the 3ʹ end of the gene, Eq. (12) also describes the number of nascent mRNAs accumulated at the 3ʹ end at time 2Δ*t*. After an infinite number of time intervals Δ*t*, the number of nascent mRNAs at the 3ʹ end *N*(∞) becomes

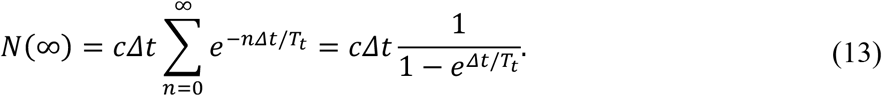

In the limit that the time interval Δ*t* goes to zero,

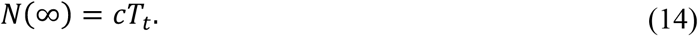

This result shows that *cT_t_* nascent mRNAs accumulate at the 3ʹ end in the steady state.

Now let us suppose transcription initiation is blocked at time *t* = 0 while in the steady state. After time *t*, the number of nascent mRNAs in the 3ʹ end becomes

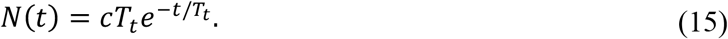

Thus, the nascent RNA density at position *x* and time *t*, *R*(*x*, *t*) can be described by the combination of the Heaviside function *H* and the Dirac delta function δ:

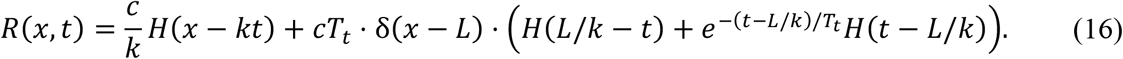

The first term in Eq. (16) is the nascent RNA density without the termination time effect. *c*/*k* indicates the number of nascent mRNAs per unit length in the occupied region. The second term describes Pol II accumulation at the 3ʹ end (*x* = *L*). The amplitude of the Dirac delta function means the number of accumulated nascent mRNAs at position *L*, which starts to decrease as the last Pol II enters the 3ʹ end (*t* = *L*/*k*). From that time, because there are no incident Pol IIs at the 3ʹ end, the amplitude of the Dirac delta function starts to decrease exponentially.

When Pol II starts to transcribe the stem–loop region [*X*_1_, *X*_2_], the intensity of pre-mRNA *I*(*x*) increases as more stem-loops are synthesized. In the post stem–loop region [*X*_2_, *L*], *I*(*x*) remains constant (Fig. 1a and 1b). Then, the intensity with the Pol II position *x* becomes

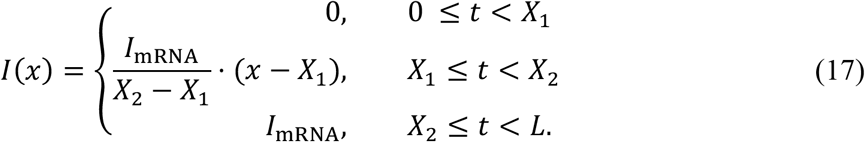

*I_mRNA_* is the intensity of a single mRNA that is fully transcribed (all 24 stem–loops). Finally, the fluorescence intensity of a transcription site at time *t*, *F*(*t*) can be derived as 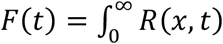. *I*(*x*)*dx*. The fluorescence intensity at *t* = 0 is

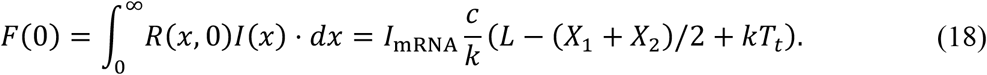

The fluorescence intensity at time *t* becomes

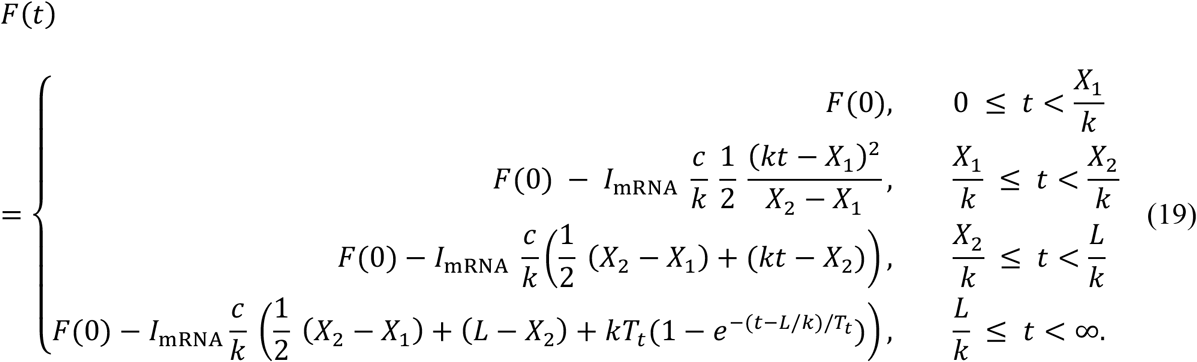

In the Eq. (19), the exponential term in the fourth interval is due to the termination time effect. A normalized fluorescence intensity *F_n_*(*t*) is

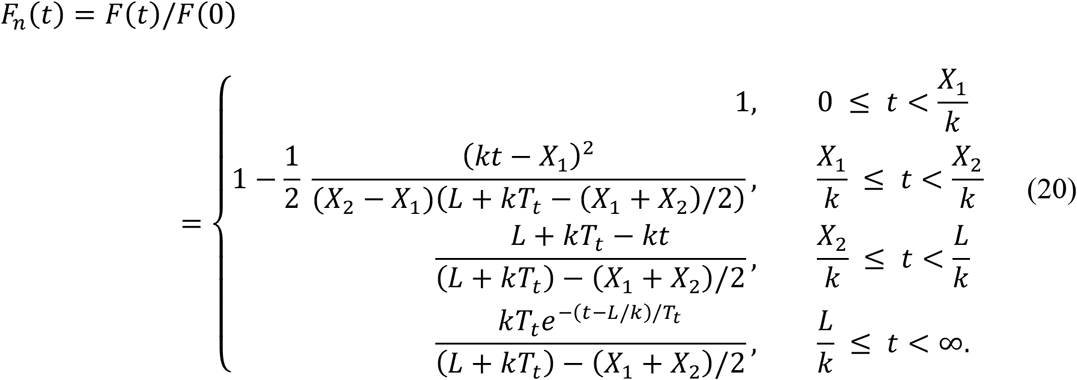

This is the model function of time-resolved transcription analysis that can be used for fitting experimental data. With the three fitting parameters, *F*(0), *k* and *T_t_*, the elongation and termination kinetics can be distinguished in single-cell experiments.

## Materials and methods

### Cell lines and reagents

Two immortalized cell lines, Actb-MBS KI^31^ MEFs and Arc-PBS KI^28^ MEFs, were used to image Actb and Arc mRNA, respectively. Lentiviral transduction was used to express MCP-GFP and stdPCP-stdGFP^32^ fusion proteins in Actb-MBS and Arc-PBS MEFs, respectively. Cells were seeded in glass-bottom dishes in Dulbecco’s modified Eagle’s medium (DMEM) (11995065, Thermo Fisher Scientific, USA) supplemented with 10% fetal bovine serum (FBS) (10082147, Gibco, US), 1% penicillin/streptomycin (Pen/Strep) (15140122, Gibco, US), and 1% GlutaMAX (35050061, Gibco, USA) and incubated at 37 °C and 5% CO_2_. Prior to imaging, cells were starved for 15-20 hours, and the medium was changed to Leibovitz’s L-15 medium without phenol red (21083027 Thermo Fisher Scientific, USA) supplemented with 1% Pen/Strep and 1% GlutaMAX. After placing the cells on a microscope, we stimulated them with a final concentration of 15% FBS. For time-resolved measurement, we blocked transcription initiation by adding 0.5 μM triptolide (T3652, Sigma Aldrich, USA) or 0.3 μM flavopiridol (F3055, Sigma Aldrich, USA) at 10 min after serum stimulation.

### Fluorescence microscopy

All images were taken using a wide-field fluorescence microscopy system based on an IX-83 inverted microscope (Olympus) equipped with a UAPON 150×/1.45 NA oil objective (Olympus), iXon Life 888 electron-multiplying charge-coupled device (EMCCD) camera (Andor), SOLA SE u-niR light engine (Lumencor), Chamlide TC top-stage incubator system (Live Cell Instrument), and ET-EGFP filter set (Chroma, ET470/40x, T495lpxr, ET525/50m). Live-cell imaging was performed at 37 °C, and time-lapse z-stack images were taken at an interval of every 8 s over a period of 67 min. Z-stacks were imaged at an interval of 0.5 μm for a total height of 6 μm.

### Image analysis

All z-stack images were maximum projected and bleach-corrected via an exponential bleach correction fit using Fiji^33^. Fluorescence intensity time traces of TSs were generated by using HybTrack^34^. The background level was determined by the average intensity of the period in which TS was not automatically detected by HybTrack. For each time trace, transcription ‘ON’ states were distinguished from ‘OFF’ states by two criteria: (1) the TS intensity was higher than 1.5 times the background level, and (2) the durations of both ON and OFF states were equal to or longer than 1 min.

For steady-state measurements, we calculated the autocorrelation function of the time traces in the ON-state; time traces longer than the average ON time were used for analysis. The autocorrelation function was calculated by the multitau algorithm^35^. This algorithm reduces noise in the correlation function at a long lag time. To measure the initiation rate and the total dwell time, we fitted an autocorrelation function of a single ON-time trace with Eq. (2). We used weighted least squares fitting with weights equal to the inverse standard error of the mean. The fitting parameters were the initiation rate *c* and the effective transition rate of Pol II *k_s_*.

For time-resolved measurements after inhibiting transcription initiation, we analyzed time traces showing a plateau followed by a decreasing signal. In our analysis, we assumed that the gene is fully and evenly occupied by Pol IIs before inhibition of transcription initiation; thus, the fluorescence intensity of TS cannot increase after inhibition. Nonetheless, some data exhibited increasing fluorescence intensity after treatment with triptolide; those time traces were considered not to satisfy the condition of our kinetic model. The selected intensity time traces were fit with the model function with a nonlinear least squares method. The fitting parameters were the elongation rate *k*, the termination time *T_t_*, and the normalization factor *F*(0).

## Results

### Steady-state fluctuation correlation analysis

To label Actb and Arc mRNA with green fluorescent proteins (GFPs), we used the MS2- and PP7-GFP systems, respectively. The MS2-GFP system utilizes highly specific binding between the MS2 bacteriophage capsid protein (MCP) and the MBS RNA stem–loop^36^. Constitutive expression of MCP fused with GFP (MCP-GFP) labeled all endogenous β-actin mRNA in mouse embryonic fibroblasts (MEFs) cultured from Actb-MBS KI mice in which 24 repeats of the MBS were inserted in the 3ʹ UTR of the β-actin gene^27^ (Fig. 1A). In a similar manner, endogenous Arc mRNA was visualized by expressing PP7 capsid protein (PCP) fused with GFP (PCP-GFP) in MEFs derived from Arc-PBS KI mice bearing 24 repeats of PBS in the Arc gene 3ʹ UTR^28^ (Fig. 1B). In the figure, the positions of the start and end of the stem–loop cassettes are denoted as X_1_ and X_2_, and the intensity change of a precursor mRNA (pre-mRNA) is shown as a function of the Pol II position *x* (Fig. 1a and b). The MS2 or PP7 capsid protein fused with GFP (CP-GFP) complexes carries a nuclear localization sequence (NLS), which targets the protein to the nucleus. When pre-mRNAs are transcribed, multiple CP-GFPs bind to the RNA binding sites (Fig. 1c), and the transcription site appears as a bright spot in fluorescence images (Fig. 1d, Supplementary video S1 and 2).

To image transcription sites, we starved homozygous Actb-MBS and Arc-PBS MEFs overnight and imaged the cells after adding serum-containing medium. A few minutes after serum induction, up to four transcription sites appeared in nuclei (Fig. 1d), and each transcription site showed bursting transcriptional activity (Fig. 2a). Both the Actb and Arc genes exhibited stochastic switching between active (ON) and inactive (OFF) states after serum induction. The duration of the ON state was 15.4 ± 6.6 min (means ± SD) for Actb, which was significantly longer than the 8.6 ± 3.4 min observed for Arc (*P* < 0.01, *t* test; Fig. 2b). The duration of the OFF state was 6.5 ± 3.4 min for Actb, shorter than the 9.4 ± 4.0 min for Arc (*P* < 0.01, *t* test; Fig. 2c). These results demonstrate that Actb exhibits transcriptional bursts more frequently in longer time periods than Arc.

**Fig. 2.**
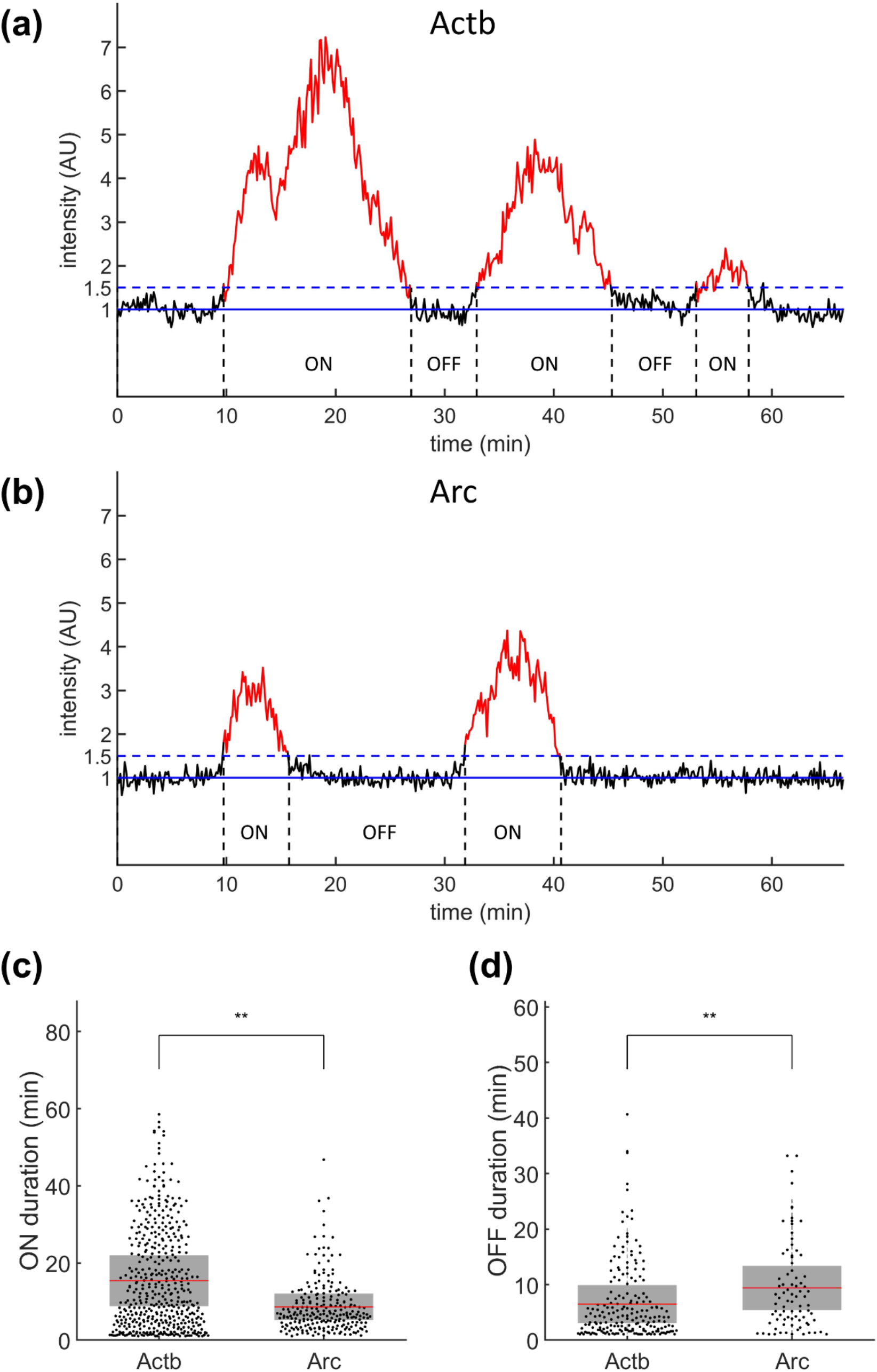
Transcription ON and OFF durations of Actb and Arc genes. (**a, b**) Representative intensity profiles of transcription sites of Actb (a) and Arc (b) genes. The solid blue lines indicate the baseline normalized by the nuclear background. The dashed blue lines indicate the threshold for transcriptional ON (red) and OFF (black) states. AU, arbitrary units. (**c, d**) ON (c) and OFF (d) durations of Actb (n = 449 for ON and 198 for OFF states) and Arc (n = 234 for ON and 83 for OFF states). The mean (red line) and standard deviation (SD, gray box) values are indicated (\**P* < 0.05 and \*\**P* < 0.01, Wilcoxon rank-sum test).

The initiation rate and total dwell time were measured from the steady-state autocorrelation function of transcription intensity trace for ON time duration. First, we characterized the general behavior of the model function in Eq. (2). The autocorrelation amplitude *G*(0) in Eq. (2) decreases when *c* increases, and *G*(0) increases when *k* increases (Fig. 3a and b). The decay time of the autocorrelation function is independent of *c* (Fig. 3a) but increases as *k* decreases (Fig. 3b).

**Fig. 3.**
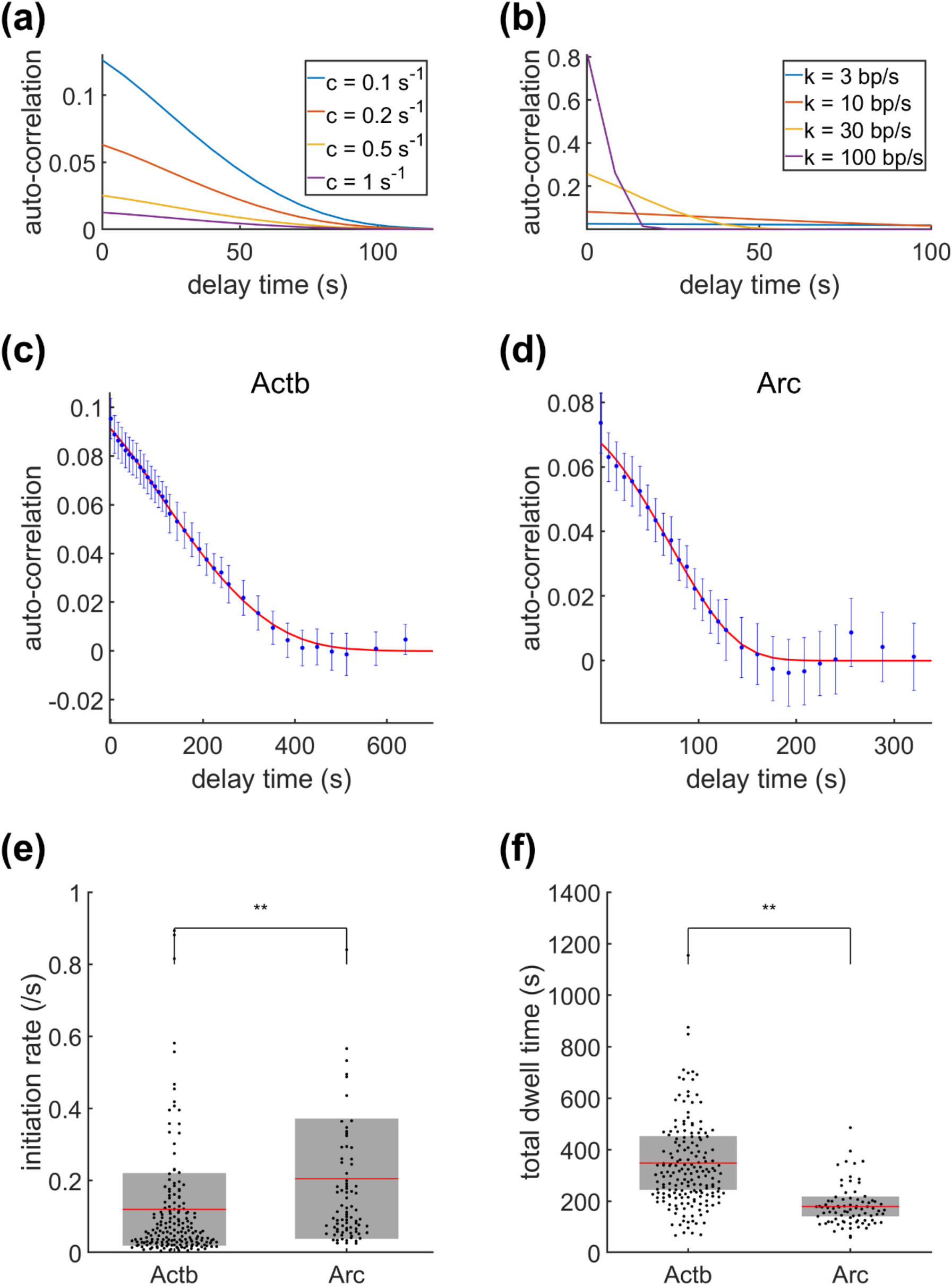
Steady-state autocorrelation analysis of transcription. (**a, b**) Autocorrelation model function behaviors with varying initiation rate *c* (a) and total dwell time *T_d_* (b). The termination time was set to *T_t_* = 0. For (a), the total dwell time was fixed at *T_d_* = 100 s, and for (b), the initiation rate was *c* = 0.1 s^−1^. (**c, d**) Representative autocorrelation data (blue dots) and the best fit curves (red lines) of Actb (c) and Arc (d) transcription. (**e, f**) The initiation rates (e) and the total dwell times (f) of Actb (n = 183 ON-time traces) and Arc (n = 82 ON-time traces). The mean (red line) and standard deviation (SD, gray box) values are indicated (\*\**P* < 0.01, Wilcoxon rank-sum test).

Representative autocorrelation curves of Actb and Arc transcription are depicted in Fig. 3c and d. The initiation rate *c* and the total dwell time *T_d_* obtained from individual ON traces are plotted in Fig. 3e and f. The average initiation rate and total dwell time were *c* = 0.12 ± 0.10 s^−1^ and *T_d_* = 348 ± 104 s for Actb and *c* = 0.21 ± 0.17 s^−1^ and *T_d_* = 179 ± 38 s for Arc.

### Time-resolved transcription analysis

Because the total dwell time *T_d_* includes the elongation and the termination time, these two dynamic factors could not be separated in steady-state measurements. Therefore, we performed time-resolved measurements to separate elongation and termination effects. We used a small molecule transcription inhibitor, triptolide (Trp) (MW 360.6), for time-resolved analysis. It has been shown that triptolide inhibits transcription initiation by preventing transcription bubble formation without significantly affecting the elongation rate^37,38^. After inhibiting transcription initiation, the region occupied by Pol IIs decreased over time in the direction of elongation (Fig. 4a). Then, the normalized intensity of a transcription site declined in 4 steps (Fig. 4b). First, the intensity remained constant in 0 ≤ *t* < *X*_1_/*k* after inhibitor addition. As the last Pol II entered the stem–loop region, the intensity decreased quadratically in the interval of *X*_1_/*k* ≤ *t* < *X*_2_/*k*. After that, the intensity decreased linearly in the time interval of *X*_2_/*k* ≤ *t* < *L*/*k* as the last Pol II entered the post stem–loop region. After time *t* = *L*/*k*, there were no elongating Pol IIs, and the intensity decreased exponentially due to the termination time effect. The model function behaviors for the different values of the termination time *T_t_* and the elongation rate *k* are shown in Fig. 4c and 4d, respectively. The increase in the termination time *T_t_* resulted in a longer decay time, though the time length of the plateau remained the same (Fig. 4c). The length of the plateau *X*_1_/*k* is solely determined by the elongation rate *k* (Fig. 4d), which denotes the speed of Pol II in the pre stem–loop region. Therefore, the elongation rate *k* and termination time *T_t_* can be acquired by fitting the normalized transcription site intensity of the experimental data with the model function in Eq. (20).

**Fig. 4.**
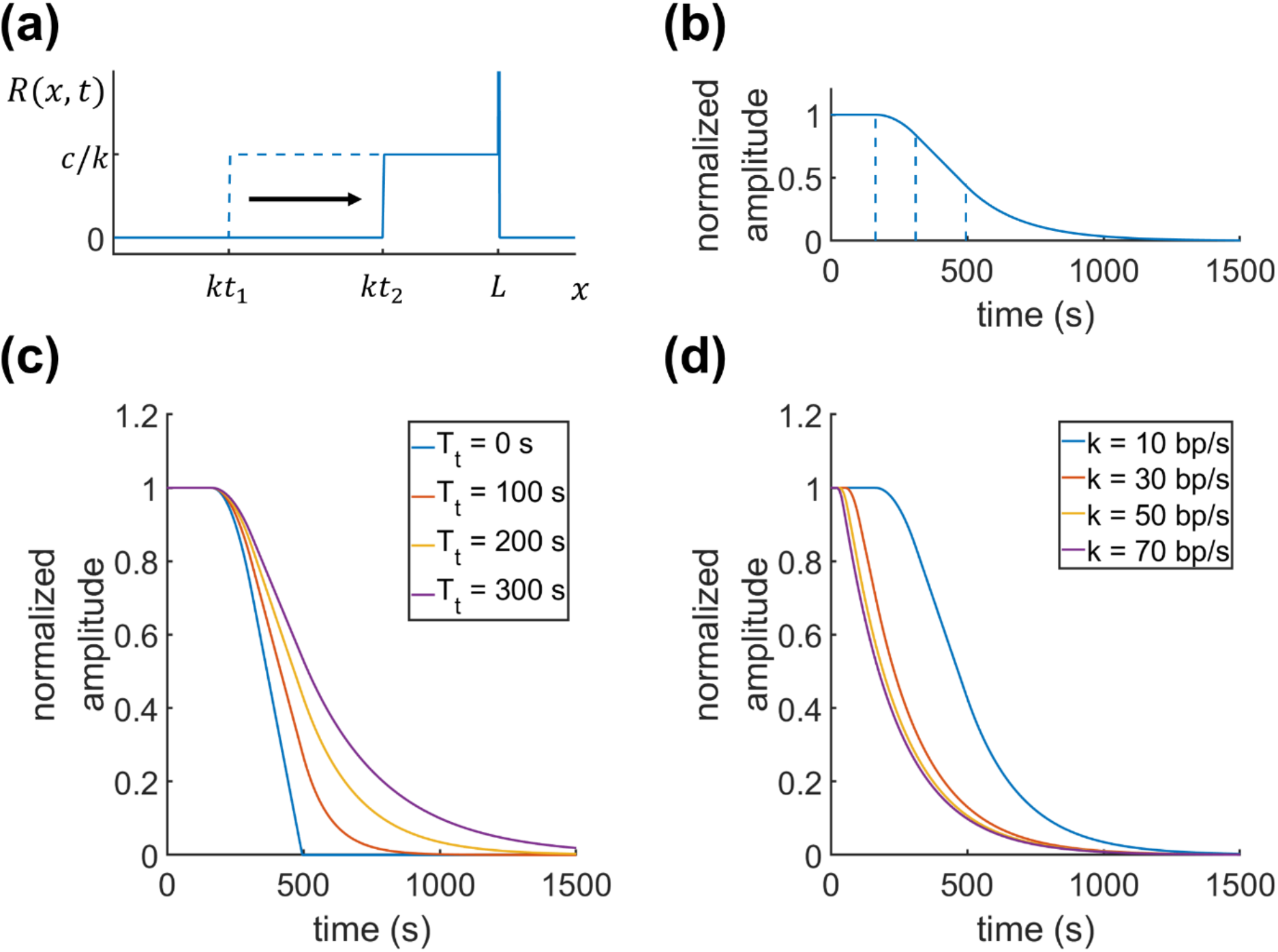
Model functional behaviors of the time-resolved transcription analysis. (**a**) The change in Pol II density distribution from *t*_1_ to *t*_2_. The transcription initiation inhibitor was added at *t* = 0. (**b**) The model function of the transcription intensity over time after adding the inhibitor. (**c, d**) The model function calculated for different termination times (c) and elongation rates (d). For (c), the elongation rate was fixed at *k* = 10 bp/s, and for (d), the termination time was *T_t_* = 200 s.

Representative data of the normalized fluorescence intensities of transcription sites in Actb-MBS and Arc-PP7 MEFs are provided in Fig. 5a and 5b, respectively. By fitting the data with Eq. (20), we obtained an average elongation rate of Actb transcription of 10.4 ± 2.9 bp/s, which was similar to the Actb transcription elongation rate measured by fluorescence in situ hybridization (FISH) in normal rat kidney (NRK) cells^39^. On the other hand, the average elongation rate of Arc was significantly different from that of Actb; the average elongation rate of Arc transcription was 29.0 ± 17.0 bp/s, almost 3-fold faster than that of Actb (Fig. 5c). Termination times also differed significantly. As illustrated in Fig. 5d, the average termination time of Actb mRNA (233 ± 59 s) was 2.7-fold longer than that of Arc mRNA (86 ± 27 s) (Fig. 5d).

**Fig. 5.**
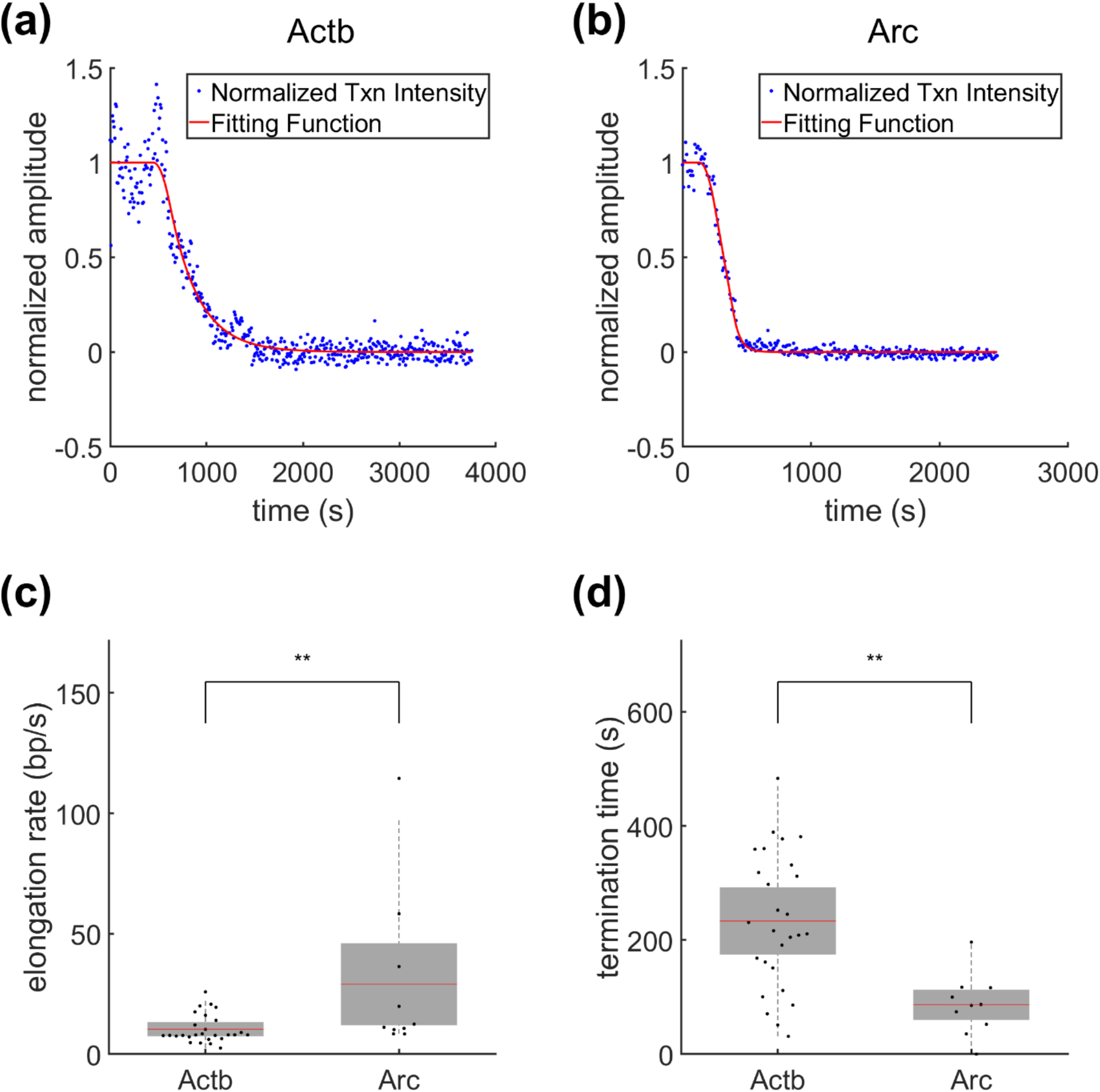
Time-resolved analysis of Actb and Arc transcription. (**a, b**) Representative transcription intensity data (blue dots) and the best fit curves (red lines) of Actb (a) and Arc (b) after the addition of triptolide at *t* = 0. (**c, d**) The elongation rates (c) and termination times (d) of Actb (n = 27 transcription sites) and Arc (n = 10 transcription sites). The mean (red line) and standard deviation (SD, gray box) values are indicated (\*\**P* < 0.01, Wilcoxon rank-sum test).

To validate our measurements, we used another transcription inhibitor, flavopiridol (FP) (MW 438.30), which is known to inhibit cyclin-dependent kinase 9 (Cdk9) in the positive transcription elongation factor b (P-TEFb) complex^40^. This P-TEFb inhibitor FP prohibits Pol II release from the promoter proximal pausing to the productive elongation phase, whereas Trp inhibits new Pol II initiation^8^. Although the mechanisms of inhibition by Trp and FP are different, they result in a similar physical state that inhibits transcription initiation. Using FP with the same procedure as in the Trp experiment, we obtained elongation rate and termination time consistent with the Trp results. We measured the elongation rates of Actb and Arc transcription from the FP experiment as 13.3 ± 5.6 and 29.0 ± 15.3 bp/s, respectively, similar to those from the Trp experiment (Fig. 6a and b). The termination times obtained from the FP experiments were 223 ± 59 and 60 ± 25 s for Actb and Arc transcription, respectively. These values were also consistent with the Trp results (Fig. 6c and d). The consistency between the two independent experiments using Trp and FP supports that our time-resolved analysis approach can measure transcriptional elongation and termination kinetics without significant toxicity effect by the drugs. We determined the average elongation rate and termination time of each gene by pooling the single-allele data set from the Trp and FP experiments (Table 1).

**Table 1.** Transcription parameters measured in this study.

|  | Actb | Arc | Unit |
| --- | --- | --- | --- |
| ON duration | $15.4 \pm 6.6$ | $8.6 \pm 3.4$ | min |
| OFF duration | $6.5 \pm 3.4$ | $9.4 \pm 4.0$ | min |
| $c$ | $0.12 \pm 0.10$ | $0.21 \pm 0.17$ | $s^{-1}$ |
| $T_d$ | $348 \pm 104$ | $179 \pm 38$ | s |
| $T_t$ (Trp + FP) | $228 \pm 59$ | $65 \pm 25$ | s |
| $k$ (Trp + FP) | $12.0 \pm 4.6$ | $29.0 \pm 15.5$ | bp/s |
| $k_{SS}$ | $12.9 \pm 6.4$ | $29.1 \pm 5.8$ | bp/s |

**Fig. 6.**
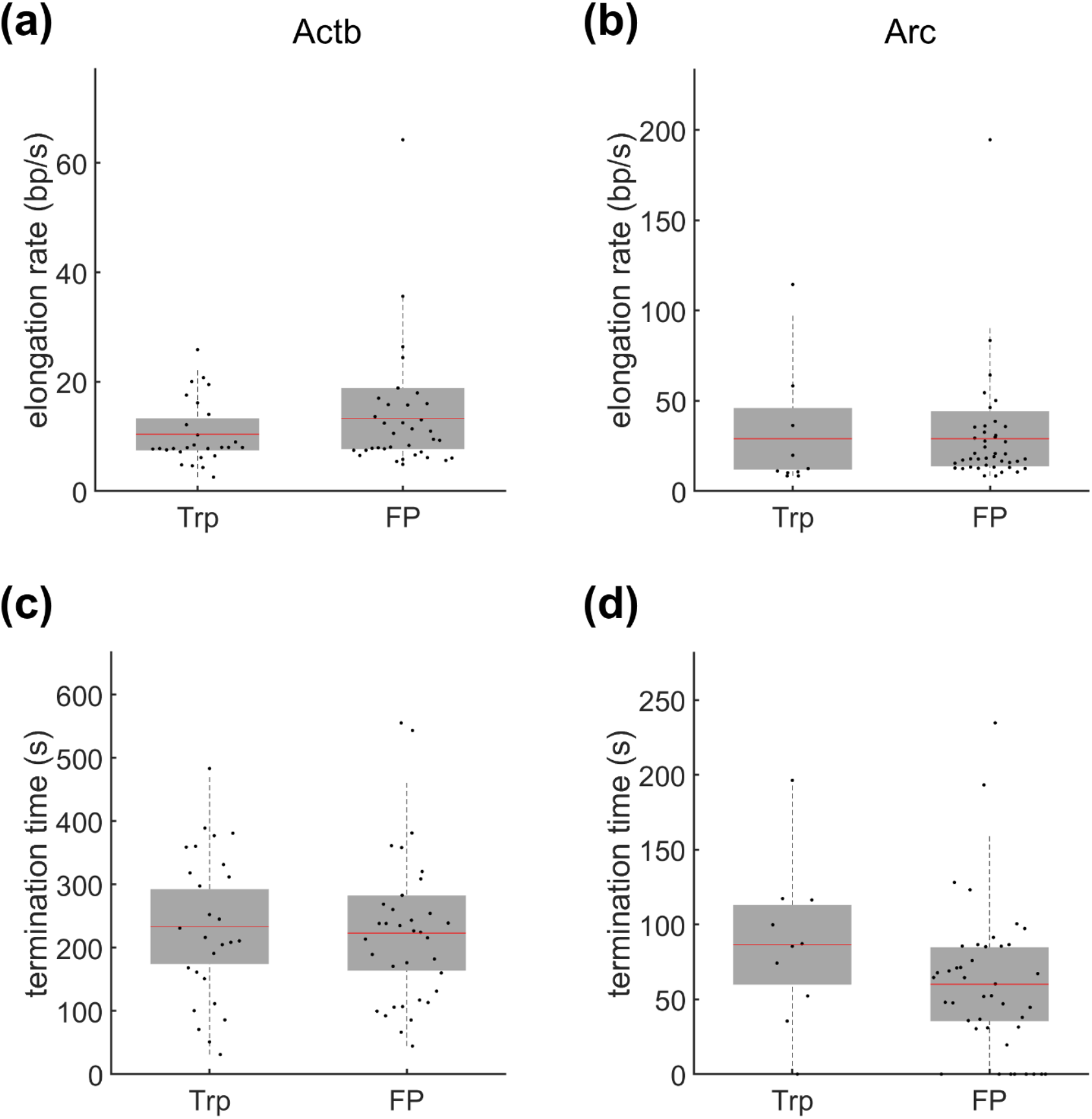
Comparison of time-resolved measurements after adding triptolide (Trp) and flavopiridol (FP). (**a, b**) The elongation rates of Actb (a) and Arc (b). The elongation rates measured after adding Trp and FP were similar for each gene. (**c, d**) The termination times of Actb (c) and Arc (d). The mean (red line) and standard deviation (SD, gray box) values are indicated. No significant difference was observed between experiments using Trp (n = 27 transcription sites for Actb and 10 for Arc) and FP (n = 35 transcription sites for Actb and 42 for Arc) (*P* > 0.05, Wilcoxon rank-sum test).

Next, we compared the elongation rates from the time-resolved measurements with those from the steady-state measurements. Using the termination time measured in the time-resolved analysis, the steady-state elongation time was calculated by subtracting *T_t_* from the total dwell time *T_d_*. Then, the elongation rate in the steady-state measurement becomes

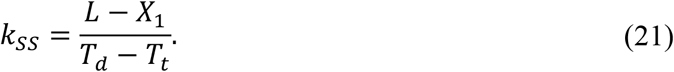

The average steady-state elongation rates were 12.9 ± 6.4 bp/s for Actb and 29.1 ± 5.8 bp/s for Arc transcription. These elongation rates depict the speed of Pol II in the fluorescently labeled regions, while the elongation rates determined by the time-resolved analysis, *k*, represent the speed in the pre stem–loop regions. These two different rates, *k_ss_* and *k*, were quite similar. Comparing the elongation rates from steady-state and time-resolved models, we affirmed that the inhibitors did not influence the elongation dynamics.

### Error estimation by simulations

Using simulations, we estimated the errors in the time-resolved and steady-state measurements corresponding to the gene structures of Actb and Arc labeled with stem-loops. The input parameters of the simulations were initiation rate *c*, elongation rate *k*, and termination time *T_t_*. From Eq. (10), the probability of *x* nucleotides elongated in time Δ*t* is

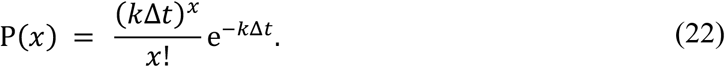

In the simulation, the time Δ*t* was 8 s, which was the same as the interval of the time-lapse imaging. Eq. (22) shows that the number of nucleotides elongated by Pol II during Δ*t* follows a Poisson distribution with a mean of *k*Δ*t*. As Pol II transcribes a stem loop, the intensity of the transcription site increases in unit brightness. After summation of all pre-mRNA intensities, the simulation generated the intensity time trace of a transcription site.

For the steady-state simulation, the total number of initiations was determined by (observation time)× (initiation rate *c*). The initiation time points were randomly generated, and the termination time was given as an exponential random number. After fitting the autocorrelation function of the simulation results, we calculated errors between the various simulation input parameters and the output results from the fitting. The error between the input and the output parameters was defined as |input − output|/input. The input total dwell time *T_d_* was calculated by *T_d_* = (*L* − *X*_1_)/*k* + *T_t_*, where *k* and *T_t_* were the input elongation rate and termination time, respectively. The mean value of the error was calculated after repeating the simulation over 100 times using the same input parameters. In Fig. 7, the color maps display the errors corresponding to the output parameters, initiation rate *c* and total dwell time *T_d_* (all values over 1 are displayed as 1). The input initiation rate *c* varied from 0.01 s^−1^ to 1 s^−1^ in the *x*-axis. The input elongation rate *k* varied from 10^0.5^ to 10^2^ bp/s in the *y*-axis. The input termination time *T_t_* is written on top in Fig. 7. Mean errors were generally greater with longer termination times *T_t_*. Additionally, a slower elongation rate *k* and a higher initiation rate *c* increased the mean error. The steady-state model function determined the average total dwell time *T_d_* of a single nascent mRNA and the average time interval 1/*c* between two nascent mRNA signals. Because fluorescence signals from multiple nascent mRNAs highly overlapped in the high *c*/*k* and *T_t_* limits, it was difficult to resolve the *T_d_* and *c* of single mRNAs in these regimes.

**Fig. 7.**
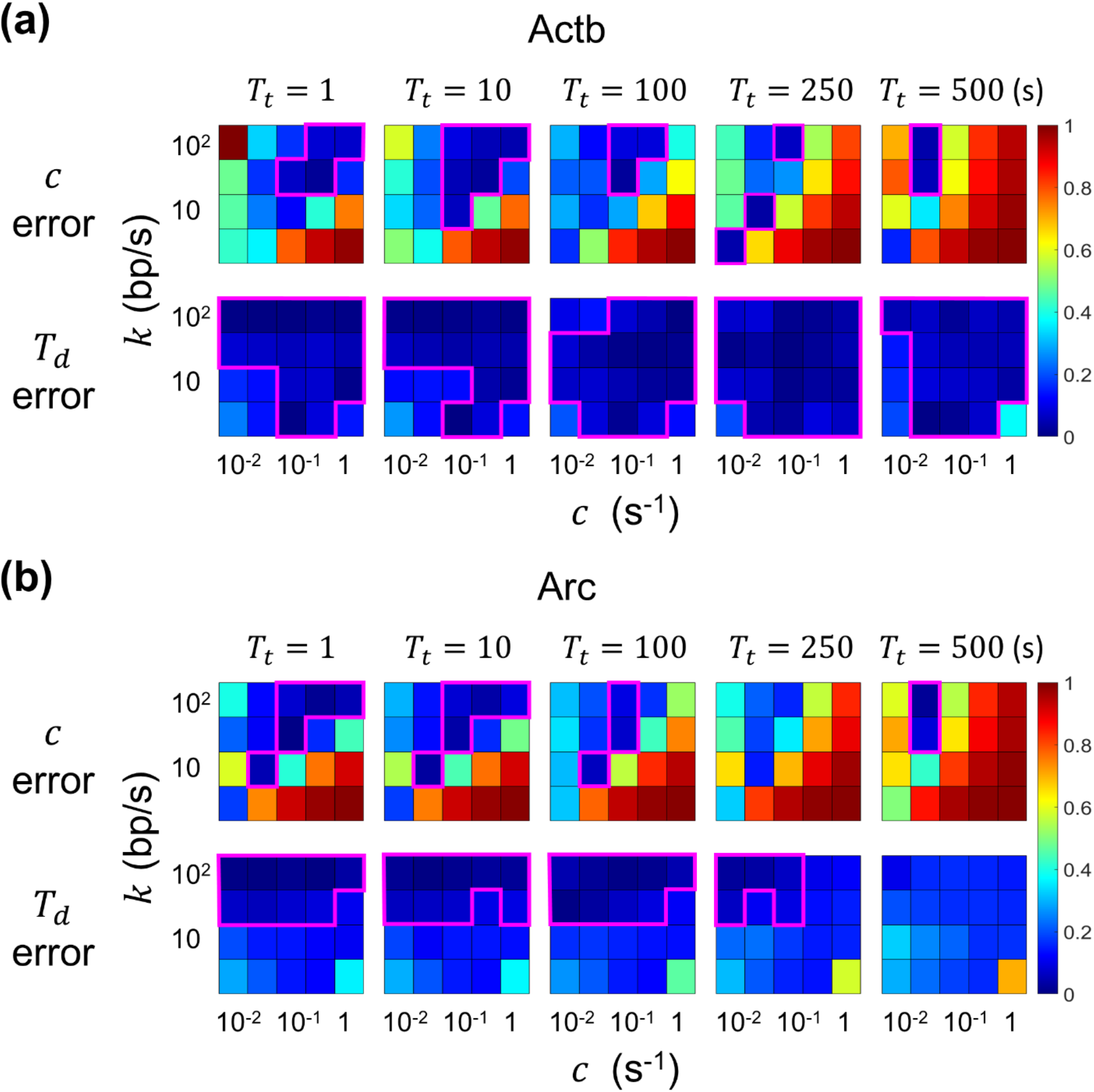
Assessment of errors in the steady-state transcription analysis by using simulations. Simulation input parameters are initiation rate *c*, elongation rate *k*, and termination time *T_t_*. (**a, b**) Mean errors of *c* and *T_d_* obtained by fitting the results from 100 simulation runs are shown in color maps for Actb (a) and Arc (b). All error values over 1 are displayed as 1. Regions with a mean error under 10% are denoted as magenta lines.

For the simulation of the time-resolved analysis, *cL*/*k* Pol IIs were distributed at random positions on the gene, and *cT_t_* Pol IIs were placed at position *L*. The termination time was given as an exponential random value, and there was no newly initiated Pol II after *t* = 0. Fig. 8 shows the errors in the elongation rate *k* and the termination time *T_t_* obtained by fitting the simulation results with Eq. (20). The error behavior of the time-resolved analysis is opposite to that of the steady-state analysis. The mean errors generally became greater in shorter termination times *T_t_* and lower initiation rates *c*. Because the assumption of uniformly distributed Pol IIs was not valid in the low *c* limits, the time-resolved analysis is not appropriate in this realm.

**Fig. 8.**
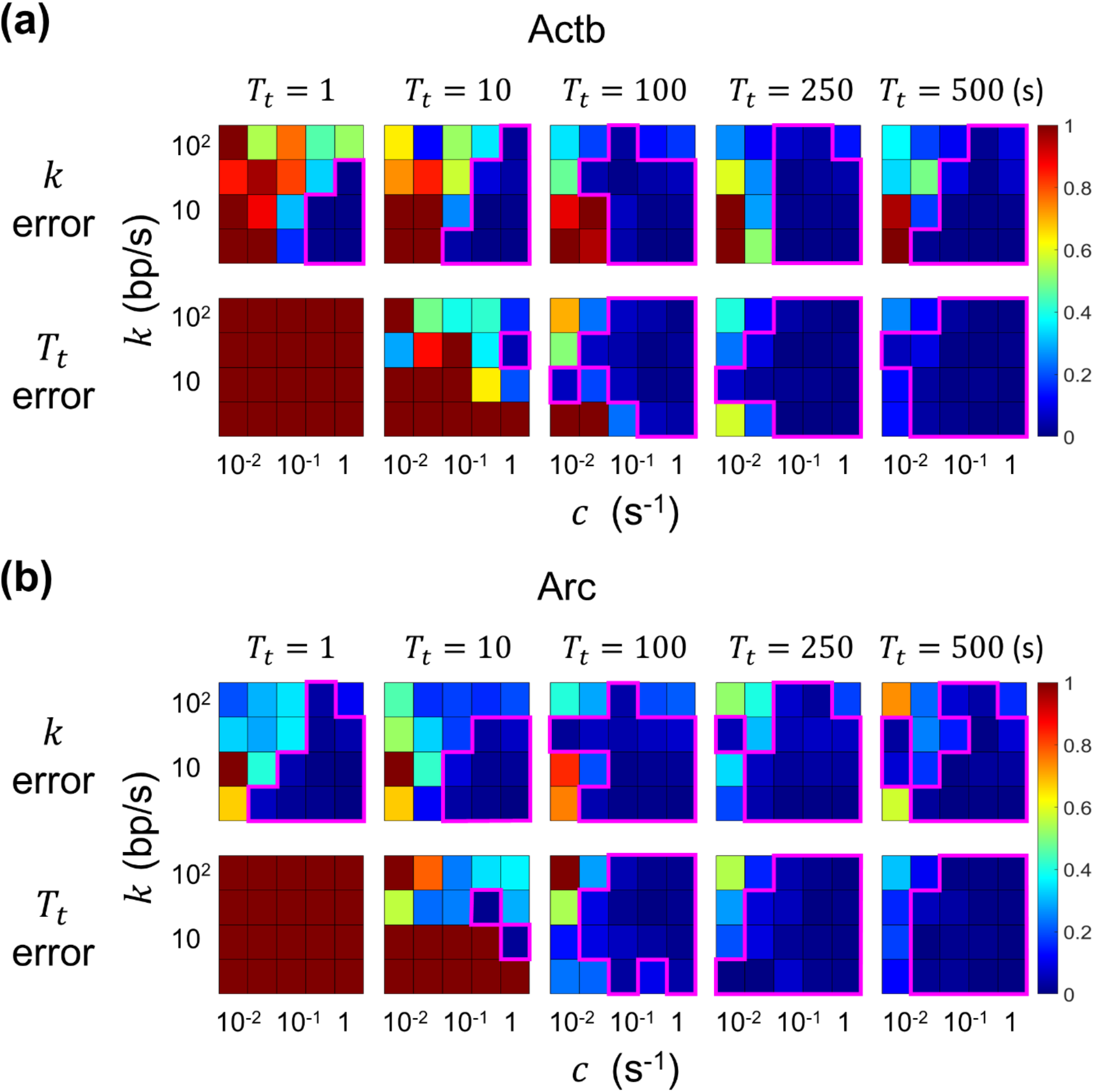
Assessment of errors in the time-resolved transcription analysis by using simulations. Simulation input parameters are initiation rate *c*, elongation rate *k*, and termination time *T_t_*. (**a, b**) Mean errors of *k* and *T_t_* obtained by fitting the results from 100 simulation runs are shown in color maps for Actb (a) and Arc (b). All error values over 1 are displayed as 1. Regions with a mean error under 10% are denoted as magenta lines.

The areas inside the magenta lines in Figs. 7 and 8 indicate the regimes in which the mean errors are under 10%. The valid region of the steady-state model function is distributed on the upper and left sides of the color map, and the time-resolved model’s valid region is on the lower and right sides. The errors in the steady-state analysis generally increased for a longer termination time, whereas the error behavior of the time-resolved analysis was the opposite. Therefore, the two analysis methods are complementary to each other. The time-resolved analysis of our Actb and Arc transcription data yielded the elongation rate *k* and the termination time *T_t_* within the valid region of less than 10% error. The steady-state analysis yielded the total dwell time *T_d_* within the 10% error region, but the initiation rate *c* of Actb was in the region with 30~50% error. These results support the overall validity of the elongation rate and termination time measured by our time-resolved method and indicate the error range of the initiation rate in the steady-state analysis.

## Discussion

In this study, we investigated the transcription kinetics of the Actb and Arc genes by using two analysis methods for live-cell imaging. Because transcription initiation, elongation, and termination dynamic factors cannot be measured separately in the steady state, we used Trp and FP to inhibit transcription initiation and performed time-resolved measurements. For simplicity, we assumed that Pol IIs accumulate at the end of the gene for transcription termination and described it as a Dirac delta function in the Pol II density distribution *R*(*x*, *t*). By fitting our data with this mathematical model, we obtained the elongation rate and the termination time separately, and the elongation rates of the Actb and Arc genes from the time-resolved measurement were consistent with those from the steady-state measurement. To identify the region in which model function fittings are valid, we checked the error behaviors of the steady-state and time-resolved models with various input parameters. The error behaviors of the two analysis methods were opposite, though the elongation rates and the termination times of Actb and Arc measured in this study were well within the valid regions of both analysis methods.

We obtained significantly different elongation rates and termination times for Actb and Arc transcription. There are many factors that affect the elongation rate of Pol II, such as exon density, nucleosome structure, and CpG content^8,11^. Previous studies have revealed that Pol II pauses or slows down at intron-exon junctions^8,9,11^ and that exon density has the greatest effect on elongation rates among the factors mentioned above^8^. The gene lengths of Actb and Arc are similar (Actb: 3640 bp, Arc: 3490 bp), but Actb has 6 exons and Arc has 3. Thus, the exon density of Actb is about two times higher (Actb: 1.6 exons/kb, Arc: 0.9 exons/kb), which explains why the elongation rate of Actb is much slower than that of Arc.

Furthermore, the termination time of Actb was 3.6-fold longer than that of Arc. There are several different types of transcription termination in mammalian cells^4^, and termination of human Actb transcription occurs by the Pol II pause-dependent termination type. This Pol II pausing occurs at the G-rich pause sequence downstream of the poly(A) signal (PAS)^41^. In the human Actb gene, nascent transcripts hybridize to the antisense DNA strand and displace the sense DNA strand, forming a structure called an R-loop^42^. The R-loop is formed prevalently over G-rich pause sites behind the elongation complex and induces Pol II pausing. Senataxin, a DNA-RNA helicase, resolves R-loops to allow the 5ʹ-3ʹ exonuclease Xrn2 to degrade the nascent RNA fragment from the poly(A) cleavage site^42^. Similar to human Actb, mouse Actb also has a G-rich region 450 nucleotides downstream of the PAS; hence, transcription of mouse Actb is likely to be terminated by the R-loop-dependent pausing mechanism. We determined the termination time, which was defined as the time from reaching the transcription end site (TES) to the release of mRNA, to be approximately 230 s for mouse Actb. These data provide us with information on the timescale of transcription termination by R-loop-dependent pausing. In contrast, the mouse Arc gene does not have a G-rich region downstream of the PAS and exhibits a much shorter transcription termination time of ~65 s. Therefore, we presume that Arc transcription termination occurs via a different mechanism. The Arc gene has a viral origin and contains sequences that are related to retrotransposon Gag genes^43,44^. It is also interesting that the transcription termination time of Arc is similar to that of the HIV-1 RNA reporter gene (63.5 s) in U2OS cells^15^. Our new analysis technique to measure termination time can be useful to unravel diverse transcription termination processes.

The transcription initiation rate was not included in our time-resolved model function because we normalized the decreasing intensity by the plateau level *F*(0). If the intensity of a single mRNA *I_mRNA_* can be measured, we can calculate the initiation rate *c* as

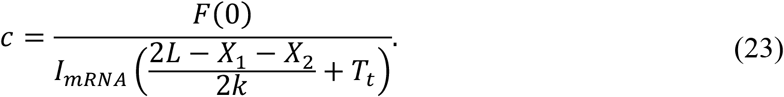

Because of the high GFP background in the nucleus, it was difficult to measure the intensities of single mRNA and transcription sites simultaneously in our live-cell experiments. We expect that background-free RNA imaging techniques such as the MS2-PP7 hybrid stem–loop system combined with split-GFP approaches^45,46^ will allow for measuring both single mRNA and transcription site intensities and extracting all three transcription kinetic rates, *c*, *k*, and *T_t_*, simultaneously in a single cell.

In addition, we expect this analysis method to be used in conjunction with RNA-targeting CRISPR-Cas systems^47–49^, through which we could analyze the transcription kinetics of any endogenous RNA without requiring genetic manipulation. Our time-resolved model function was calculated by integrating the product of the Pol II density function and the fluorescence intensity of pre-mRNA. Such model function for a different RNA tagging system can be easily derived following the same steps described herein. This analysis technique will expand our capability to investigate the transcription kinetics of endogenous genes in a highly quantitative manner.

## Data Availability

The results of experimental and simulation data are uploaded at zenodo (https://zenodo.org/record/5788345#.YbxTrtBByUl), and the source codes for analysis and simulation are uploaded at github (https://github.com/Neurobiophysics/SingleCellTxnDyn).

## Acknowledgments

This work was supported by the Creative-Pioneering Researchers Program through Seoul National University, Howard Hughes Medical Institute (HHMI)-Wellcome International Research Scholar Award from the Wellcome Trust [208468/Z/17/Z], and the Basic Science Research Program through the National Research Foundation of Korea (NRF) grant funded by the Korean government (2020R1A2C2007285).

