## Supplementary Video captions for "Time-resolved analysis of transcription kinetics in single live mammalian cells"

**This PDF file includes:**

Captions for Supplementary Videos S1 and S2

**Other supplementary materials for this manuscript:**

Supplementary Videos S1 and S2

### Supplementary Video Captions

**Video S1.** Time-lapse movie of Actb transcription after serum stimulation in the Actb-MBS MEF shown in Fig. 1d. The movie is played at 30 frame per second (fps).

**Video S2.** Time-lapse movie of Arc transcription after serum stimulation in the Arc-PBS MEF shown in Fig. 1d. The movie is played at 30 fps.
